# Multi-scale computational study of the mechanical regulation of cell mitotic rounding in epithelia

**DOI:** 10.1101/037820

**Authors:** Ali Nematbakhsh, Wenzhao Sun, Pavel A. Brodskiy, Aboutaleb Amiri, Cody Narciso, Zhiliang Xu, Jeremiah J. Zartman, Mark S Alber

**Affiliations:** Department of Applied and Computational Mathematics and Statistics, University of Notre Dame, Notre Dame, IN 46556, USA; Department of Chemical and Biomolecular Engineering, University of Notre Dame, Notre Dame, IN 46556, USA; Department of Physics, University of Notre Dame, Notre Dame, IN 46556, USA; Department of Mathematics, University of California, Riverside, CA 92521, USA

## Abstract

Mitotic rounding during cell division is critical for preventing daughter cells from inheriting an abnormal number of chromosomes, a condition that occurs frequently in cancer cells. Cells must significantly expand their apical area and transition from a polygonal to circular apical shape to achieve robust mitotic rounding in epithelial tissues, which is where most cancers initiate. However, how cells mechanically regulate robust mitotic rounding within packed tissues is unknown. Here, we analyze mitotic rounding using a newly developed multi-scale subcellular element computational model that is calibrated using experimental data. Novel biologically relevant features of the model include separate representations of the sub-cellular components including the apical membrane and cytoplasm of the cell at the tissue scale level as well as detailed description of cell properties during mitotic rounding. Regression analysis of predictive model simulation results reveals the relative contributions of osmotic pressure, cell-cell adhesion and cortical stiffness to mitotic rounding. Mitotic area expansion is largely driven by regulation of cytoplasmic pressure. Surprisingly, mitotic shape roundness within physiological ranges is most sensitive to variation in cell-cell adhesivity and stiffness. An understanding of how perturbed mechanical properties impact mitotic rounding has important potential implications on, amongst others, how tumors progressively become more genetically unstable due to increased chromosomal aneuploidy and more aggressive.

**Author Summary:** Mitotic rounding (MR) during cell division which is critical for the robust segregation of chromosomes into daughter cells, plays important roles in tissue growth and morphogenesis, and is frequently perturbed in cancerous cells. Mechanisms of MR have been investigated in individual cultured cells, but mechanisms regulating MR in tissues are still poorly understood. We developed and calibrated an advanced subcellular element-based computational model called Epi-Scale that enables quantitative testing of hypothesized mechanisms governing epithelial cell behavior within the developing tissue microenvironment. Regression analysis of predictive model simulation results reveals the relative contributions of osmotic pressure, cell-cell adhesion and cortical stiffness to mitotic rounding and establishes a novel mechanism for ensuring robustness in mitotic rounding within densely packed epithelia.

## Introduction

Epithelia are tissues composed of tightly adherent cells that provide barriers between internal cells of organs and the environment and are one of the four basic tissue types in the human body [1–3] (Fig. 1). Epithelial expansion driven by cell proliferation is a key feature throughout development, and it also occurs in hyperplasia, a precursor to cancer. Cell divisions during development must occur robustly, as mis-segregation of chromosomes leads to severe genetic abnormalities such as aneuploidy [4]. Over 90% of all human tumors are derived from epithelia [5], and the accumulation of genetic errors during cell division can lead to all of the hallmarks of cancer [6]. In tissues, mitotic cells must become sufficiently round to avoid the mis-segregation of chromosomes all the while still remaining connected with their neighbors [7]. A deeper understanding of the biophysical mechanisms governing the behavior of mitotic cells in epithelia will result in a better understanding of many diseases including cancer.

**Fig. 1.**
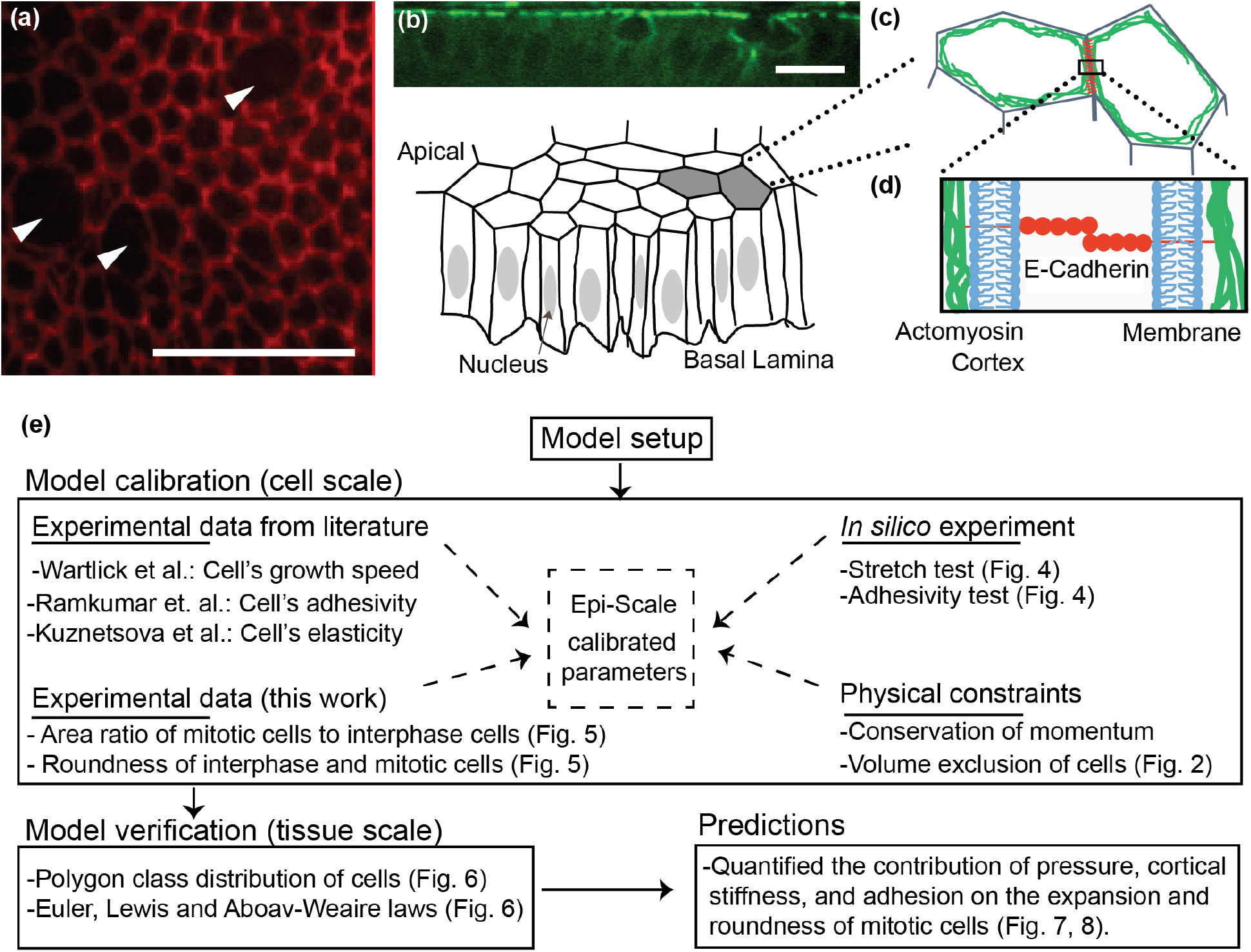
Epithelial mechanics and workflow outline. (a) Apical surface of epithelial cells within the *Drosophila* wing imaginal disc that are marked by E-cadherin tagged with fluorescent GFP (DEcadherin:: GFP). Multiple cells within the displayed region are undergoing mitotic rounding with a noticeable decrease in fluorescent intensities of E-Cadherin. (b) Experimental image of crosssection of wing disc marking levels of actomyosin (Myosin II::GFP) and cartoon abstraction of epithelial cells, which are polarized with apical and basal sides. Actomyosin and mechanical forces during mitotic rounding are primarily localized near the apical surface. (c) At the molecular scale, the boundary between cells consists of a lipid bilayer membrane for each cell, E-cadherin molecules that bind to each other through homophilic interactions, and adaptor proteins that connect the adhesion complexes to an underlying actomyosin cortex that provides tensile forces along the rim of apical areas of cells. Arrows indicate mitotic cells. Scale bars are 10 micrometers. (e) The graphical workflow of the computational modeling setup, calibration, verification and predictions.

Epithelial cells entering mitosis rapidly undergo structural changes that result in the apical area of the cell becoming larger and rounder, in a process known as mitotic rounding (MR) [8, 9]. MR occurs in detached cells, cells adherent to a substrate as well as in epithelial cells within tissues [10–12]. MR in epithelia coincides with an increased polymerization of actomyosin at the cell cortex, which results in an increase in cortical stiffness [4,11]. Simultaneously, the intracellular pressure increases [11], and cells partially reduce adhesion to their neighbors and the substrate [4].

However, the roles of cell-cell adhesion, cell stiffness, and intracellular pressure during mitotic rounding are not fully resolved in cultured cells, and even less is known in the tissue context [13]. For example, Stewart et al. [11] indicates that both pressure and the actin-myosin cortex are important for mitotic swelling while Zlotek-Zlotkiewics et al. [14] observe that the actin-myosin cortex is not involved in mitotic swelling. Further, it is technically challenging to modulate the mechanical properties of individual mitotic cells in tissues with small perturbations that do not “break” the system. Thus, this gap-in-knowledge is currently extremely hard to address experimentally.

Recently, computational modeling coupled with experimentation has become a powerful tool for identifying the biophysical mechanisms governing organogenesis [15–20]. MR is investigated in this paper by using a novel multi-scale sub-cellular element model (SEM) called Epi-Scale that simulates epithelial cells in growing tissues. New biologically relevant features of the model include: i) separate representations of the apical membrane and cytoplasm, as well as cell-cell interactions at the tissue scale; ii) a systematic calibration of the model parameters to provide accurate biological simulations of cell division and tissue growth; and iii) a detailed description of cell properties during mitotic rounding.

We used multi-scale model simulations and response surface methodology (multiple linear regression) [21,22] to investigate the extent to which a mitotic cell regulates cell-cell adhesion, cortical stiffness, and internal pressure and to analyze the impacts of changes in these cell properties on cross-sectional areas of mitotic cells at the apical surface as well as their roundness. The quantitative analysis of model simulations demonstrated that increasing cytoplasmic pressure was the main driver of the increase of mitotic cell’s apical area, which was balanced by both cortical stiffness and cell adhesivity. Increased cortical stiffness and decreased adhesion was shown to promote cell roundness. Surprisingly, within the range of experimentally observed MR values, the relative roundness of cells was not sensitive to small perturbations in cytoplasmic pressure. Understanding how perturbed mechanical properties such as cytoplasmic pressure, cell-cell adhesion and cortical stiffness impact mitotic rounding have important implications on, amongst others, how tumors progressively become more genetically unstable (chromosomal aneuploidy) and more aggressive.

The paper is organized as follows. The Methods section describes modeling background and new model description. The Results section provides details of calibration of single cell model parameters using quantitative experimental data. Calibrated model simulations are shown to predict emergent properties of epithelial topology without requiring further calibration using tissue-level properties. The model is then used to quantify the relative impacts of cell-cell adhesion, membrane stiffness and intracellular pressure on MR using two separate criteria: apical area and apical roundness. The paper ends with the Discussion section, which puts predictions of the model in more general biological context. It also describes future extensions of the computational model environment for simulating epithelial tissue mechanics in greater biological detail.

## Methods

### Modeling background

Multiple computational approaches have been utilized to model various aspects of epithelial tissue dynamics, each with its particular focus and applications (see, amongst others, reviews [15,23–27]). For example, the cellular Potts modeling (CPM) approach has been used successfully to take into account cell adhesivity for studying cell aggregation as well as cell morphogenesis (see, amongst others, [28–31]). Finite element models (FEMs) and models based on solving Naiver-Stokes equations have also been implemented to investigate cell growth and division [32–35]. Vertex based models (VBM) provided an efficient and fast approach to study regulation of cell topology, tissue-size regulation, tissue morphogenesis, and the role of cell contractility in determining tissue curvature [23,36–41]. In VBMs, cellular shapes are defined by the shared vertices of neighboring cells and edges between them.

The Subcellular Elements Model (SEM), developed initially by Newman’s group [42] for simulating multi-cellular systems to encompass multiple length scales, has been now adopted by many groups as a general computational modeling approach. A particular advantage of the SEM approach is that it can provide local representations of mechanical properties of individual cells which can be directly related to the experimental data [43]. Each cell in a SEM consists of a set of nodes representing a coarse-grained representation of subcellular components of biological cells. Node-node interactions are represented by energy potentials. SEMs have been extended to predict how mechanical forces generated by cells are redistributed in a tissue and for studying tissue rheology, blood clot deformation, and cell-cell signaling [44–46]. For example, a SEM model with GPU implementation was used to compare multiple mechanisms governing the formation of stratified layers of the epidermis [19] as well as mechanisms governing intestinal crypt homeostasis [47]. Jamali et al. [48] also developed an SEM model to represent the membrane and nucleus of the cell by nodes connected by overdamped springs. Gardiner et al. [49] described a SEM with locally-defined mechanical properties. Christely et al. [45] have developed an efficient computational implementation of the SEM simulating role of Notch signaling in cell growth and division, on GPU clusters to decrease computational time. A SEM model was also used to study aspects of epithelial cell mechanics without making assumptions about cell shapes [50].

### Multi-component computational model of epithelia

We describe in this section novel multi-scale SEM computational platform called Epi-Scale which simulates the growth of flat epithelial monolayers. Model simulations focus on representing two-dimensional (2D) planar cell shapes near the apical surfaces of cells of the *Drosophila* wing imaginal disc, which is popular model to study the biophysics and genetics of epithelial tissue growth. The 2D planar model is a common simplifying approximation that was used in many previous models of wing disc growth [18,38,39,51,52]. In particular, it is reasonable to use a 2D model for studying many epithelial processes in the *Drosophila* wing disc pouch because it consists of a single layer of cells and the essential structural components of those cells, including E-cadherins and actomyosin, are concentrated on the apical surface of the epithelia (Fig. 1a-d). E-cadherin is responsible for adhesion between two neighboring cells, and actomyosin, which is concentrated near the apical surface, drives cell contractility. The nucleus and most of the cytoplasm are pushed up to the apical surface during cell division. Using a 2D approximation also allows us to model a large number of cells with high resolution and with special attention to mechanical cell properties. The future development of the Epi-Scale simulation platform implemented on GPU clusters, will also enable 3D simulations with reasonable computational costs.

In what follows, we first describe different types of the sub-cellular nodes that are used to simulate each cell, and the interactions between them. Then, the equations of motion of each subcellular element are provided. Finally, approaches for modeling cell’s growth, transition to mitotic phase, and division are described. The workflow of the model is shown in Fig. 1e.

### Sub-cellular elements

Epi-Scale represents individual cells as collections of two types of interacting subcellular elements: internal nodes and membrane nodes (Fig. 2). The internal nodes account for the cytoplasm of the cell, and the membrane nodes represent both plasma membrane and associated contractile actomyosin cortex. The internal and membrane nodes are placed on a 2D plane, representing the apical surface of epithelia.

**Fig. 2.**
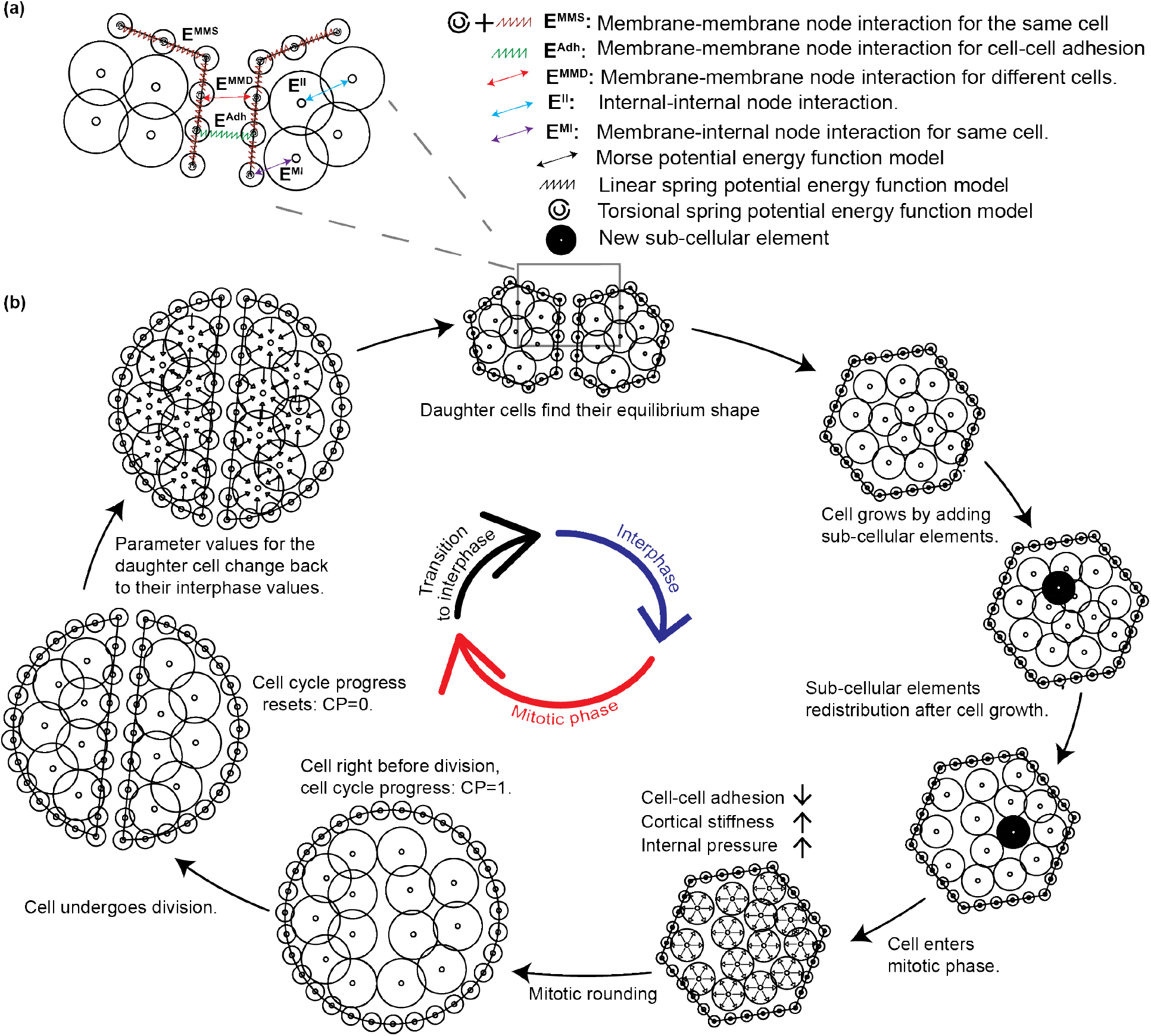
Diagram of the underlying physical basis of model simulations. (a) Intracellular and intercellular interactions between different elements of the model. Symbols and notations are indicated in the legend. (b) Implementation of the simulation of cell cycle in the model.

Interactions between internal and membrane nodes are modeled using potential energy functions as shown in Fig. 2a [45,53]. Combined interactions between pairs of internal nodes (*E^II^*) represent the cytoplasmic pressure of a cell. Combined interactions between internal nodes and membrane nodes of the same cell (*E^MI^*) represent the pressure from cytoplasm to the membrane. Interactions between membrane nodes of the same cell (*E^MMS^*) are used to model the cortical stiffness. Cell-cell adhesion (*E^Adh^*) is modeled by combining pairwise interactions between nodes of the membranes of two neighboring cells. *E^MMD^* is a repulsive Morse potential function between membrane nodes of neighboring cells that prevents membranes of adjacent cells from overlapping. Epi-Scale utilizes spring and Morse energy potential functions to simulate the interactions between subcellular elements. Linear and torsional springs are represented by energy functions *E^MMS^* and *E^Adh^* [34,54], while Morse potential functions are used in energy function *E^MI^*, *E^II^*, and *E^MMD^* [46] (see Fig. 2). The Morse potential consists of two terms, generating short-range repulsive and long-range attractive forces [42]. For example, the following expression is a Morse potential function used in *E^MI^* to represent an interaction between internal node *i* and membrane node *j:*

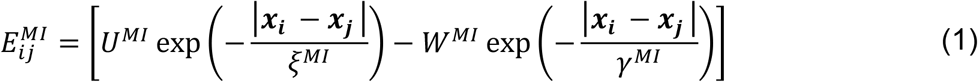

where *U^MI^*, *W^MI^*, *ξ^MI^*, and *γ^MI^* are Morse parameters. The same form of the potential with different sets of parameters is used for *E^II^* and *E^MMD^* (Table 2). These potential functions govern the motion of internal and membrane nodes inside the cells resulting in the deformation and rearrangement of cells within the tissue. A complete list of all potential functions used in the Epi-Scale to model mechanical properties of cells and epithelial tissue and description of their biological relevance are provided in Table 1 and described in SI-S.2.

**Table 1:** Potential energy functions in the Epi-Scale model.

| Potential function | Type of potential function | Biological concept |
| --- | --- | --- |
| Internal-internal nodes ($E^{II}$) | Morse | Internal pressure |
| Membrane-internal nodes ($E^{MI}$) | Morse | Keeps the cytoplasm inside the cell and applies pressure from the cell's cytoplasm to the cell's membrane |
| Membrane-membrane nodes of neighboring cells ($E^{MMD}$) | Morse | Volume exclusion of the cells (Fig. 2) |
| Membrane-membrane nodes of neighboring cells ($E^{adh}$) | Linear spring | Adhesion between neighboring cells |
| Membrane-membrane nodes of the same cell ($E^{MMS}$) | Linear and torsional spring | Membrane and cortex stiffness of the cell |

**Table 2:** Energy function parameters.

| Parameter | Interphase | Mitotic phase | Values during interphase & mitosis | Source or calibration section |
| --- | --- | --- | --- | --- |
| $E^{II}$ | $U_{inter}^{II}$ | $U_{mit}^{II}$ | 0.49 & 21.75 $nN \cdot \mu m$ | Fig. 5 and SI-S3.4 |
| | $W_{inter}^{II}$ | $W_{mit}^{II}$ | 0.15 & 6.71 $nN \cdot \mu m$ | |
| | $\xi_{inter}^{II}$ | $\xi_{mit}^{II}$ | 0.31 & 0.58 $\mu m$ | |
| | $\gamma_{inter}^{II}$ | $\gamma_{mit}^{II}$ | 1.25 & 1.34 $\mu m$ | |
| | $L_{inter}^{II}$ | $L_{mit}^{II}$ | 1.56 $\mu m$ & 3.12 $\mu m$ | |
| $E^{MI}$ | $U_{inter}^{MI}$ | $U_{mit}^{MI}$ | 0.78 & 4.36 $nN \cdot \mu m$ | Fig. 5 and SI-S3.4 |
| | $\xi_{inter}^{MI}$ | $\xi_{mit}^{MI}$ | 0.13 & 0.27 $\mu m$ | |
| | $L_{inter}^{MI}$ | $L_{mit}^{MI}$ | 1.56 $\mu m$ & 3.12 $\mu m$ | |
| $E^{MMD}$ | $U_{inter}^{MMD}$ | $U_{mit}^{MMD}$ | 3.9 $nN \cdot \mu m$ | Volume exclusion of the cells (Fig. 2) |
| | $W_{inter}^{MMD}$ | $W_{mit}^{MMD}$ | 3.9 $nN \cdot \mu m$ | |
| | $\xi_{inter}^{MMD}$ | $\xi_{mit}^{MMD}$ | 0.13 $\mu m$ | |
| | $\gamma_{inter}^{MMD}$ | $\gamma_{mit}^{MMD}$ | 1.6 $\mu m$ | |
| | $L_{inter}^{MMD}$ | $L_{mit}^{MMD}$ | 0.78 $\mu m$ | |
| $E^{adh}$ | $k_{inter}^{Adh}$ | $k_{mit}^{Adh}$ | 20 & 8.0 $nN/\mu m$ | [61,69,70] and Fig 5 |
| | $L_{max}^{Adh}$ | $L_{max}^{Adh}$ | 0.40 $\mu m$ | |
| | $L_{min}^{Adh}$ | $L_{min}^{Adh}$ | 0.062 $\mu m$ | |
| $E^{MMS}$ | $k_{inter}^{Stiff}$ | $k_{mit}^{Stiff}$ | 200 & 450 $nN/\mu m$ | [67,68] and Fig. 5 |
| | $L_{inter}^{Stiff}$ | $L_{mit}^{Stiff}$ | 0.060 & 0.13 $\mu m$ | |
| | $k_{inter}^{Tor}$ | $k_{mit}^{Tor}$ | 6.0 & 7.0 $nN \cdot \mu m/rad$ | |

\*\* Other Morse parameters are equal to zero. Mitotic values represent the central point in the CCD experimental design (Fig. 7a).

### Equations of motion of individual nodes

Displacement of each internal or membrane node is calculated at each moment in time based on the potential energy functions. The model assumes that nodes are in an overdamped regime [20,38,53] so that inertia forces acting on the nodes can be neglected. This leads to the following equations of motion describing movements of internal and membrane nodes, respectively:

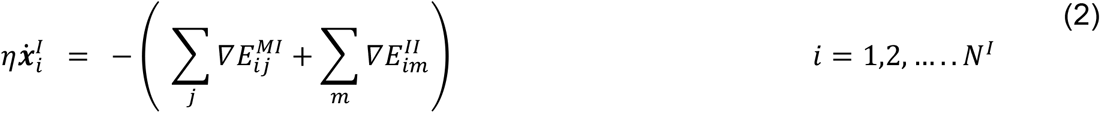

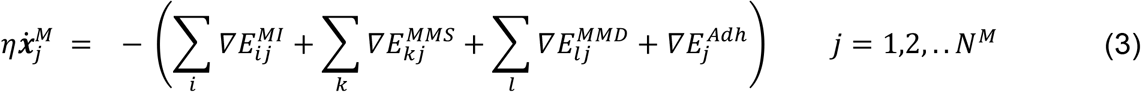

where *η* is the damping coefficient, *x_i_^I^* and *x_j_^M^* are positions of internal node and membrane nodes indicated by indices *i* and *j*. *m* is the index for any internal node interacting with the internal node *i*. *k* is the index for any membrane node of the same cell interacting with the membrane node *j*. Finally, *l* is the index for any membrane node of different cell interacting with the membrane node *j*. Note that adhesion between membranes of two neighboring cells is represented as pair-wise interaction between membrane nodes. Consequently, no summation with respect to different nodes is needed in Eqn. 3. Eqn. 2 & 3 are solved at the same time for all *N^I^* internal nodes and *N^M^* membrane nodes.

Eqns. 2 and 3 are discretized in time using forward Euler method and positions of nodes *x_i_^I^* and *x_j_^M^* are incremented at discrete times as follows

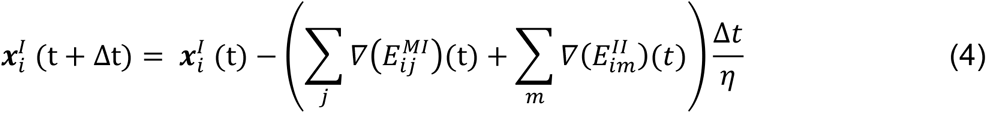

where Δ*t* is the time step size. The same discretization technique is used for the equations of motion of the membrane nodes.

Epi-Scale platform is computationally implemented on a cluster of Graphical Processing Units (GPUs). This enables us to run simulations with subcellular resolution at the micro-scale with a reasonable computational cost and to study the impact of changes in individual cell mechanical properties on the tissue development at the macro-scale. Supplementary Information (SI-S1) provides details about the simulation algorithm, GPU implementation and computational cost.

### Cell cycle

Model parameters were set based on experimental values determined from studies of *Drosophila* wing disc development, an established genetically accessible model of organ development [55]. The growth of the wing disc is spatially uniform and decreases over time [56]. The growth rate for cell *i* is modeled by an exponentially decaying function fit to the experimental data for *Drosophila* wing disc [56], with a random term representing stochastic variation among cells:

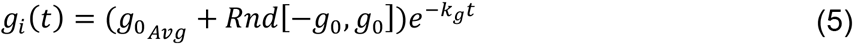

where *g_0_Avg__* is the average growth rate of cells in the beginning of the simulation and *Rnd* [−*g_0_, g_0_*] is a random number chosen using a uniform distribution in the range of [−*g_0_, g*_0_]. *k_g_* is the decay constant of the growth rate.

Cells cycle through interphase and mitosis phases in the simulation. The variable Cell Progress (*CP ϵ* [0,1]) describes progress of a cell through the cell cycle from the beginning of the interphase (*CP* = 0) to the end of the cell division (*CP* = 1). *CP* is updated based on cell growth rate as follows:

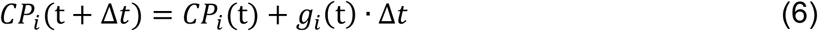

The number of internal nodes of a cell increases as the cell grows. The number of initial and final nodes can be varied based on the desired resolution of a single cell. Simulations in this work start with 20 internal nodes at CP=0 and end with 40 internal nodes at CP=1. So, an internal node is added for every 1/20 increase in CP (Fig. 2) to reach the desired 40 internal nodes at the end of cell cycle. The new internal node is randomly placed within a radius 0.2*R_c_* from the center of the cell, where *R_c_* is the radius of the cell. Epithelial cells undergoing mitosis increase their intracellular pressure by adjusting their osmolarity relative to their surroundings [57]. Additionally, the actomyosin cortex is enriched, and cellular adhesion to the substrate and to neighboring cells are downregulated [11,58–62]. Since these changes in mitotic cells occur concurrently, the relative impact on a mitotic cell cannot be easily decomposed in experiments into separable effects.

To simulate MR, parameters regulating cell-cell adhesion, cortical stiffness, and internal pressure of cells in the mitotic phase (M phase) are varied linearly from interphase parameter values to mitotic parameter values to represent the changes in cell mechanical properties during mitosis (see Table 2 and SI-S3.3) [10,11,61]. For example, *U^MI^*, Morse parameter that determines cytoplasmic pressure on the membrane of the cell (see SI-S3.4), was varied from the interphase value (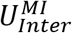) to the mitotic value (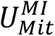), by using the following function:

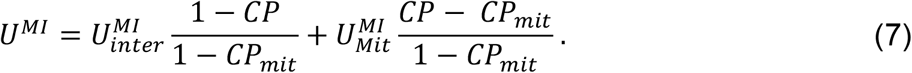

Similar linear expressions are used for representing enrichment of the actomyosin cortex and reduction in cell adhesion with neighboring cells in mitotic phase (Table 1).

Cells in the mitotic (M) phase – which lasts approximately 30 minutes – divide into two daughter cells (Fig. 2). Cytokinesis occurs when CP approaches 1 and is modeled by separating internal and membrane nodes of the mother cell into two sets representing daughter cells. The axis of division is implemented perpendicular to the cell’s longest axis, following Hertwig’s rule [63], prior to the initiation of mitotic rounding [64]. New membrane nodes are created along the cleavage plane for each daughter cell. After division, parameters for nodes of each daughter cell are set back to calibrated interphase values and *CP* is set to zero for both daughter cells.

Membrane nodes in the beginning of a simulation are arranged in a circle for each cell, and internal nodes are randomly placed within each cell (Fig. 3a). After initialization, internal nodes rapidly rearrange in every cell and cells self-organize into a polygonal network, similar to the experimentally observed cell packing geometry of epithelia (Fig. 3b). Cells in a simulation constantly grow, divide and interact with each other resulting in a detailed dynamic representation of the developing epithelial tissue (see Fig. 3c-d and movie SI-S8.1).

**Fig. 3.**
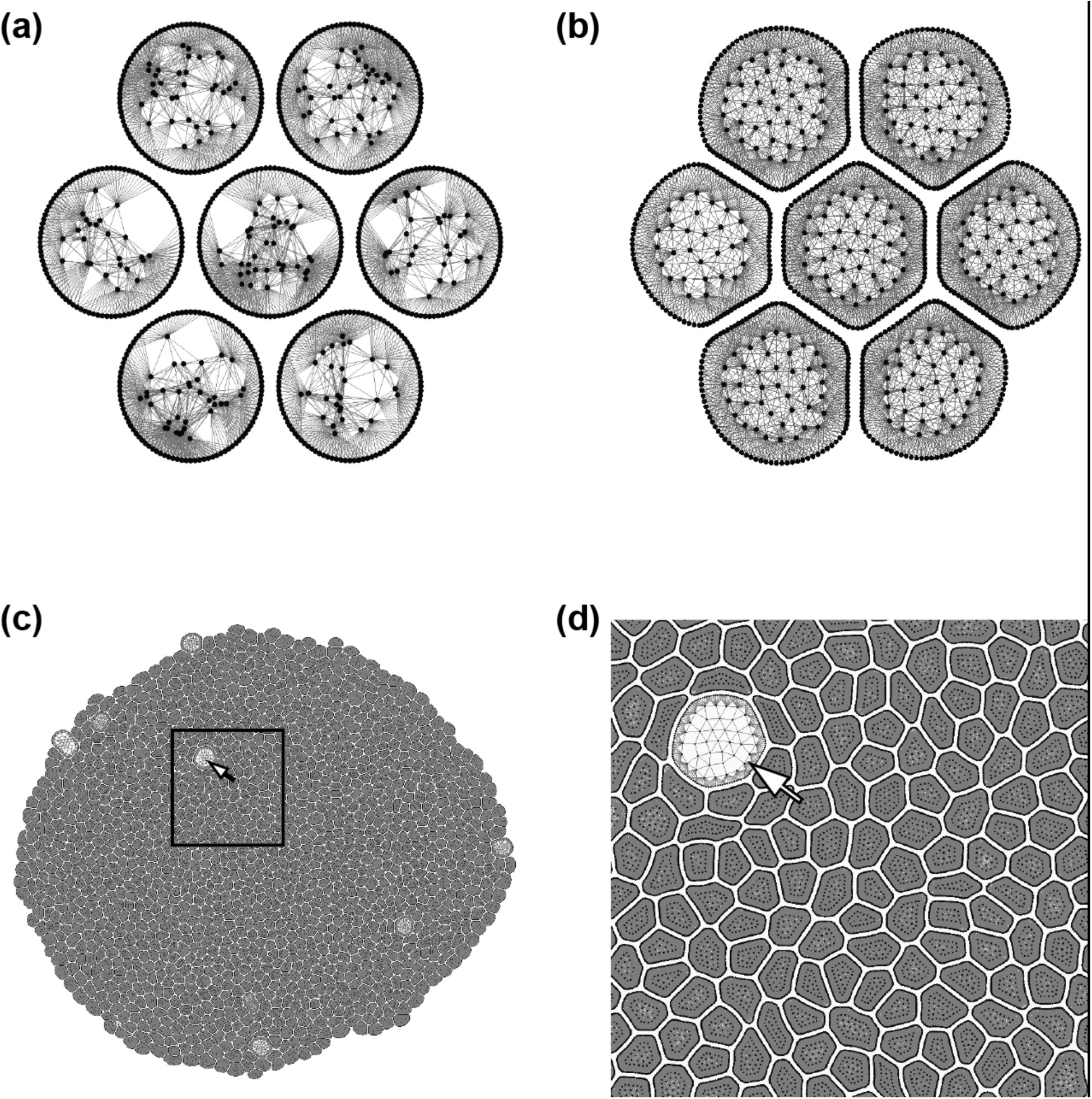
Initial conditions and sample simulation output. (a) Initial condition of a simulation with seven initially non-adherent circular cells. Each cell starts with 100 membrane elements and 20 internal elements. (b) Initial formation of an epithelial sheet after cells adhere to each other. An equilibrium distribution of internal nodes is reached for each cell. (c) Epithelial sheet after 55 hours of proliferation. (d) Enlarged view of the selected region showing different cell shapes and sizes due to interactions between cells. The large cell is undergoing mitotic rounding (MR).

## Results

### Model Calibration

Model parameters were calibrated using experimental data for the third instar *Drosophila* wing disc, which is a powerful model for studying organ formation [24,65] (Figs. 4 and 5). Experimental values for similar cell lines were used to calibrate the model parameters when experimental data for *Drosophila* wing disc were not available (Tables 1-3).

**Fig. 4.**
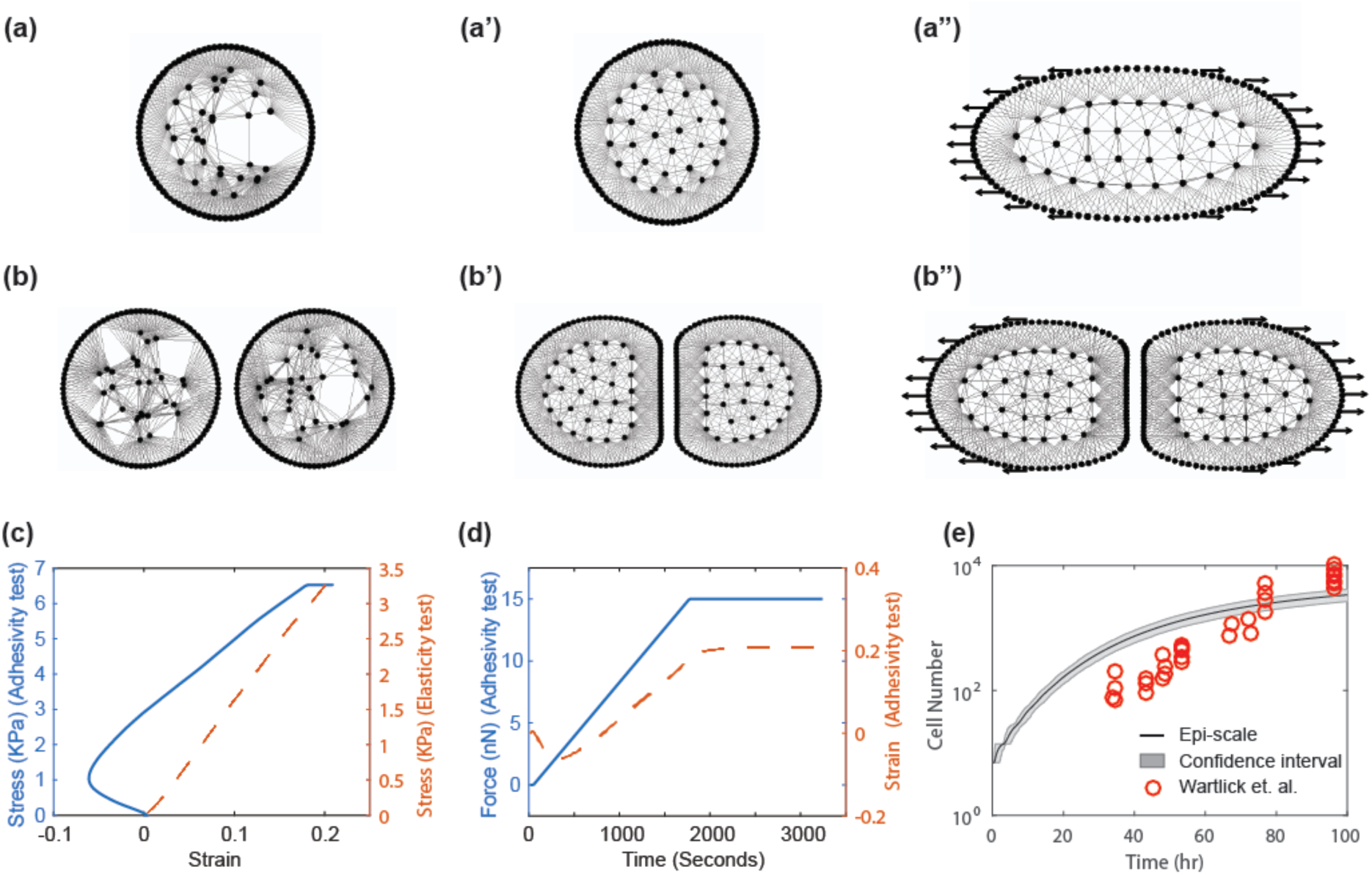
Calibration of model parameters through simulations. (a-a″) Calibration test to determine parameters for cell elasticity, analogous to experimental single cell stretching tests [66], (a) Initial condition t=0, (a') 6 minutes after simulation with no force applied, (a″) after 72 minutes cell is completely on tension (b-b″) Cell adhesivity test, analogous to experimental tests [69] for calibrating the level of cell-cell adhesion between adjacent cells. (b) Initial condition *t*=0, (b') 6 minutes after simulation begins with no force applied, (b″) after 72 minutes, 15 nN force is applied. (c) Stress versus strain for single cell calibration (red line) and stress versus strain for calibrating the level of adhesivity between the two cells (blue line) [69,70]. Initial negative strain in adhesivity test is due to strong adhesion between two cells. (d) Force and strain as a function of time for adhesivity test. (e) Tissue growth rate calibration by comparing with the experimental data by Wartlick et al. [56]. The 95% confidence interval for the growth rate results is shown in grey color.

**Fig. 5.**
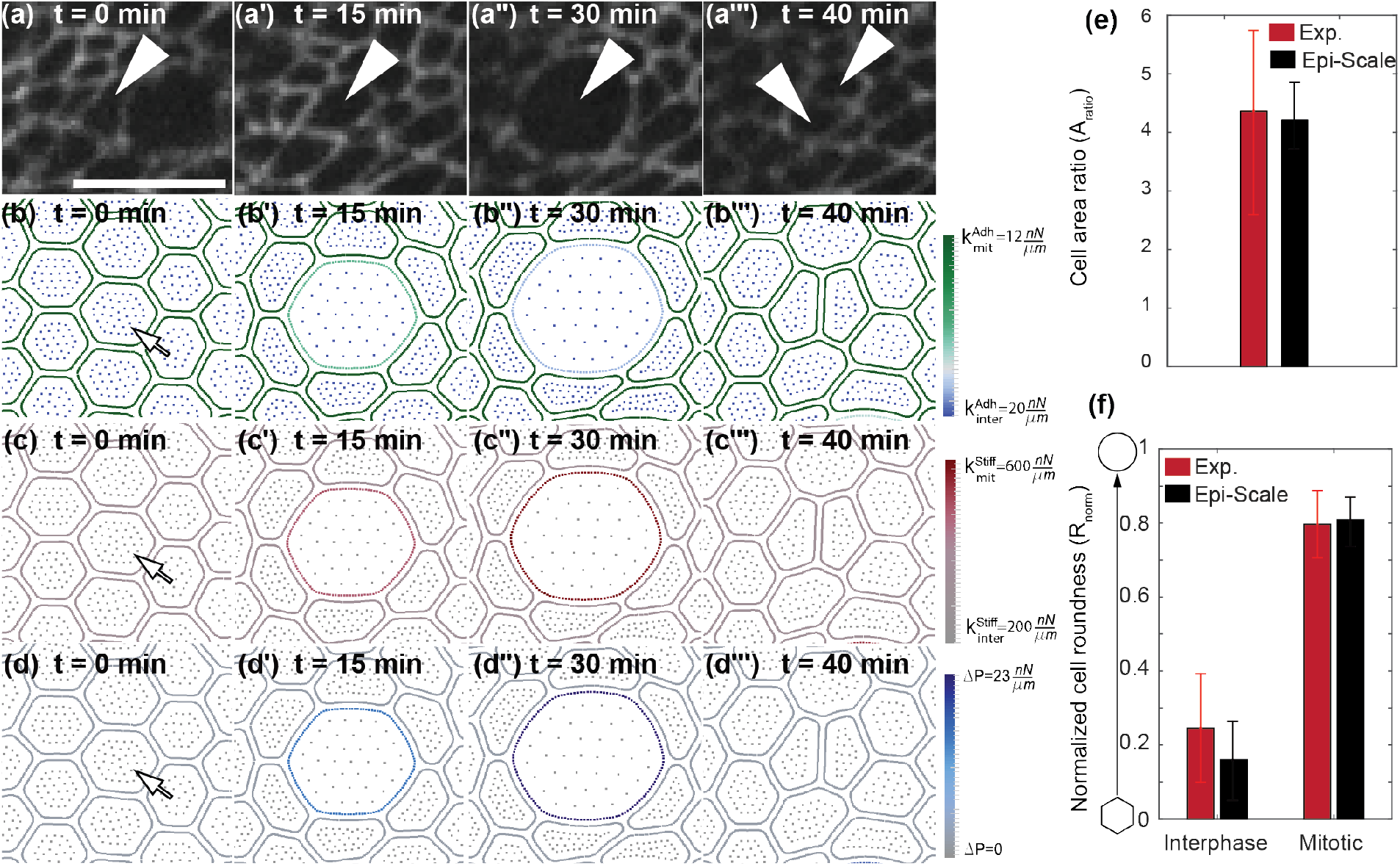
Dynamics of mitotic rounding. (a-a’”) Time-lapse confocal images of cell undergoing mitosis in the wing disc with E-Cadherin:GFP-labeled cell boundaries. Scale bar is 5 μm. Arrows indicate daughter cells. (b-d’”) Time series from Epi-Scale simulation of a cell undergoing mitosis and division with illustration of: (b) adhesive spring stiffness, (c) cortical spring stiffness, and (d) internal pressure, respected to their interphase values. (e-f) Comparison of size and roundness of mitotic cells with experimental data for the *Drosophila* wing disc. Arrow represents mitotic cell in (b-d). A t-test comparing the means of computational simulations and experiments result in p=0.72 for cell area ratio and p=0.76 for normalized roundness of mitotic cells.

**Table 3:**
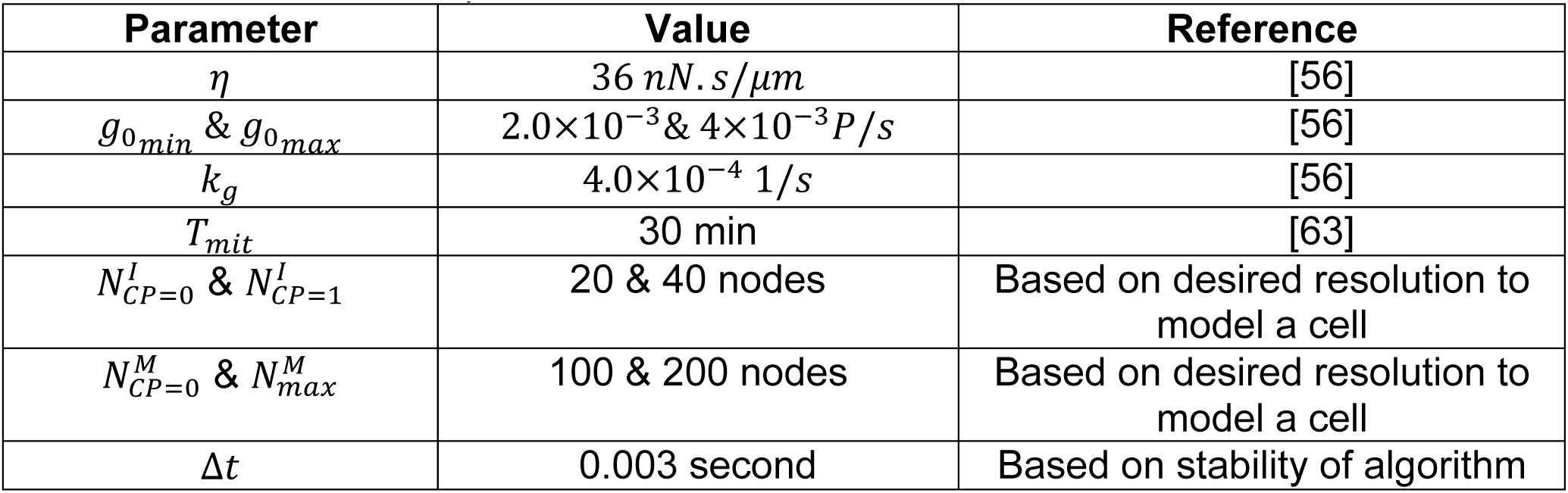
Implementation parameters

| Parameter | Value | Reference |
| --- | --- | --- |
| $\eta$ | $36 \text{ nN} \cdot \text{s} / \mu\text{m}$ | [56] |
| $g_{0\min}$ & $g_{0\max}$ | $2.0 \times 10^{-3}$ & $4 \times 10^{-3} \text{ P/s}$ | [56] |
| $k_g$ | $4.0 \times 10^{-4} \text{ 1/s}$ | [56] |
| $T_{mit}$ | 30 min | [63] |
| $N_{CP=0}^I$ & $N_{CP=1}^I$ | 20 & 40 nodes | Based on desired resolution to model a cell |
| $N_{CP=0}^M$ & $N_{max}^M$ | 100 & 200 nodes | Based on desired resolution to model a cell |
| $\Delta t$ | 0.003 second | Based on stability of algorithm |

The mechanical stiffness of the actomyosin cortex (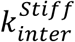) was calibrated using the modulus of elasticity (*E*) of a single cell [66]. *E* was experimentally obtained by applying forces to opposite sides of a cell and measuring cell deformation [67,68]. This experiment was reproduced in the Epi-Scale model simulation by applying a linearly increasing force to membrane nodes on both sides of a simulated cell and calculating cell’s deformation (Figure 4a-a”). The slope of the graph of the stress versus strain (Figure 4c) provides elasticity of the cell. The elasticity of a single cell is calibrated by adjusting linear stiffness of springs representing interactions between membrane nodes of a single cell. We have chosen value of the (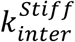) so that that *E* = 19 *kPa*, which is within the biological range of 10 − 55 *kPa* measured for epithelial cells [67,68].

The cell-cell adhesive force (***F**_adh_*) is experimentally determined by measuring the force needed to detach two adhered cells from each other. This experiment is reproduced *in silico* by applying forces to membrane nodes on either side of two adhered cells, and measuring the force needed to separate them (Fig. 4b-b”). The strength of the cell-cell adhesion for the *Drosophila* epithelium has not been measured yet. 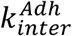 was calibrated so that 10 *nN/μm* was required to detach two adhered cells from each other, based on published data for S180 cells transfected to express E-Cadherin [69] and data from epithelial MDCK cells, which have adherens junctions similar to those along the apical surface of the *Drosophila* epithelium [70] (Fig. 4d). (More details about cell-cell adhesion calibration are provided in SI-S.5.)

Cells in the wing disc have spatially-uniform growth-rates that slow down as the tissue approaches its final size [56]. The growth rate in the Epi-Scale model described by Eqn. (5) was calibrated (Table 3) so that the number of cells in the model simulations matched experimental data for the wing disc pouch [56] (Fig. 4e).

During mitosis, apical cell area and roundness increase compared to their interphase values (Fig. 5a-a’”). This correlates with an observed increase in the cell’s internal pressure, cortical stiffness and decrease of intercellular adhesion marked by noticeable reduction in E-Cadherin. To simulate this, parameters of *E^II^* and *E^MI^*, which can be changed to increase the cytoplasmic pressure (ΔP) (SI-S3.4), cortical stiffness (*k^Stlff^*), and adhesivity (*k^Adh^*) are varied from their interphase values to their mitotic values (Fig. 5b-d). Values were selected such that the ratio of mitotic cell area to interphase cell area (*A_mit_/A_inter_*) and cell roundness (*R*) were calibrated to data collected on mitotic cells from the wing disc (Fig. 5e-f). The methods for calculating of area, roundness, and pressure are described in SI-S3.2, S3.3, and S3.4, respectively.

### Tissue topology emerges from cell self-organization driven by cellular mechanics

After calibration of the model parameters at the cellular scale, validation simulations were run to determine whether the cellular-scale calibration was sufficient to recapitulate expected topological properties of the tissue (Fig. 6a) [71,72]. One metric for tissue topology is the distribution of cell neighbor numbers, or polygon class distribution. The polygon class distribution in Epi-Scale simulations approaches to steady state after 35 hours (Fig. 6b). This steady state distribution matches the distributions observed in experiments with the wing disc and other epithelial systems [43] (Fig. 6c) as well as obtained using other computational models such as vertex based model [38]. We also confirmed that simulations recapitulate experimental observations [63] that cells entering mitosis on average gain a cell bond, increasing the number of neighbors by one (Fig 6d, inset). Further, we investigated the effects of varying cellular modulus of elasticity on the polygon class distribution in the range reported values of epithelial cells (10 − 55 *kPa*) [67]. The results show that the polygon class distribution is insensitive to the changes in the elasticity values (Fig 6e). This is a reasonable result since the polygon class distribution is strongly conserved among a wide range of epithelial tissues (Figure 6c). Therefore, cellular modulus of elasticity observed for the range of epithelial cells does not impact the polygon class distribution.

We further verified that the polygon class distribution of the simulated tissue satisfies three laws describing topological relationships: Euler’s law, Lewis law, and Aboav-Weaire Law. Euler’s law states that cells forming a packed sheet should be hexagonal on average [72,73]. The Lewis law states that cells with more neighbors should have larger normalized area [73]. The Aboav-Weaire law indicates that the average polygon class of neighbors of each cell decreases as the cell’s polygon class increases [74]. Simulation results obtained using calibrated model, show the average side of cells to be equal to 5.98 for interphase and mitotic cells, 5.80 for interphase cells, and 6.49 for mitotic cells. Model simulations also satisfy two other laws as shown in Fig. 6d when interphase cells are counted.

**Fig. 6.**
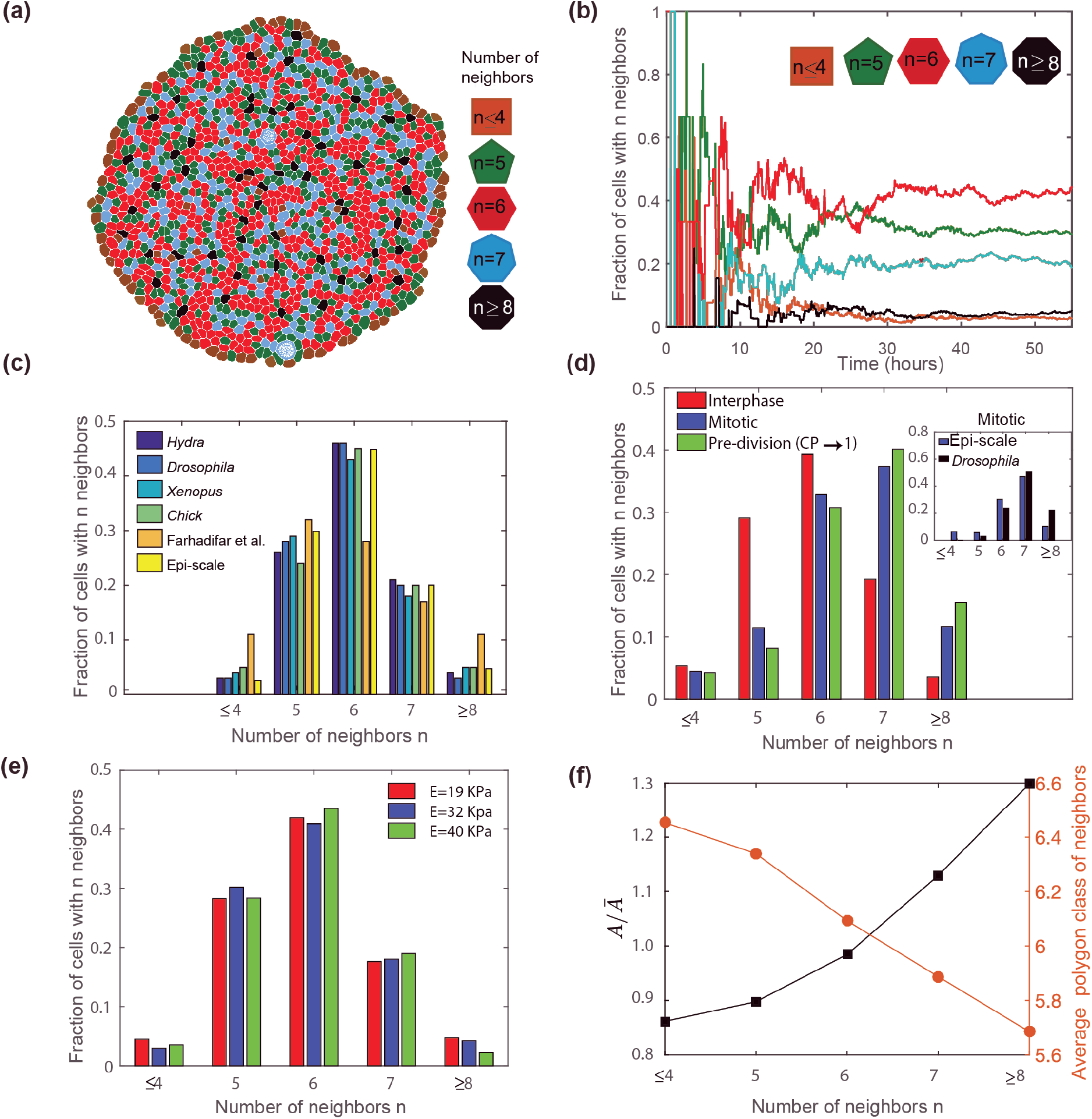
Emergence of tissue-level statics from model simulations. (a) Sample simulation output showing cells with different numbers of neighbors as different colors (b) Simulations initiated from seven cells reaches steady-state polygon-class distribution after approximately 35 hours of cell proliferation. (c) Comparison of polygon class distributions obtained by Epi-Scale model with various biological systems (data extracted from [79]) and a vertex based model by Farhadifar et al. [38]. (e) Polygon class distribution of cells at different stages of growth, and comparison of mitotic cells distribution with *Drosophila* wing disc experimental data [63]. (e) Polygon class distribution of cells at different level of cell’s elasticity. The results do not show sensitivity in the range of reported elasticity of epithelial cells [67]. (f) Average relative area (*A/Ᾱ*), and average polygon class of neighboring cells verifying that simulation results satisfy Lewis law and Aboav-Weaire law. *A* is the apical area of cell and *Ᾱ* is the average apical area of the population of cells.

### Impacts of adhesion, stiffness, and cytoplasmic pressure on mitotic rounding

The Epi-Scale model is suitable for generating and testing hypotheses regarding mechanical mechanisms of MR because it is capable of representing non-polygonal shapes of cells. Simulations were conducted to predict the relative contributions of different cell properties to the relative area ratio (*A_mit_/A_inter_*) and normalized roundness (*R-norm*) of mitotic cells as calculated in SI-S3.2 and SI-S3.3. Parameter values were selected in a three-level full factorial design (FFD) to investigate the relationships between the mitotic parameters of the model, (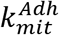, 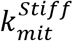, *and* Δ*P*) and mitotic rounding (*A_ratio_* and *R_norm_*) (Fig. 7a, SI-S9.1). A regression model was fit to the results of the FFD, and showed that for large changes, ΔP was the primary regulator of *A_ratio_*, and 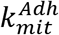 and 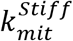 were the primary regulators of *R_norm_* (Fig. 7c-d).

**Fig. 7.**
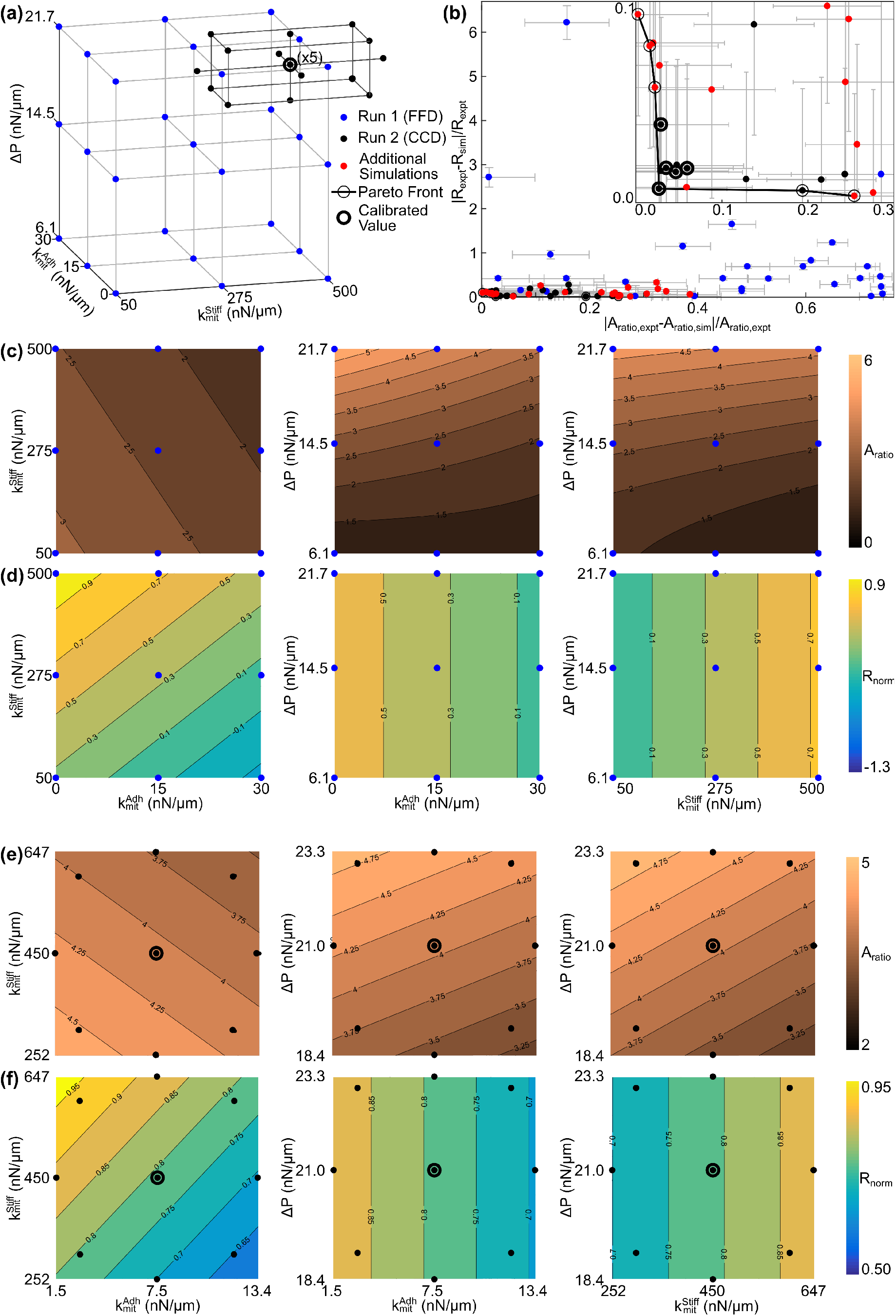
Response surface method analysis of mechanical properties on regulating mitotic expansion and mitotic rounding. (a) Schematic of initial full factorial design (FFD) for exploring parameter space, and subsequent central composite design (CCD) for developing the response surface models shown in (c, d). (b) Pareto front indicating computational model parameter values with lowest difference with experimental data for area ratio and normalized roundness. The parameter range defined by the CCD (Run 2) spans parameter variation where the error between experiments and simulations is within the propagated uncertainty of measurements and simulations. Error bars are the standard error of means of the normalized deviation between experiments and simulations. (c-d) Contour plots for FFD experiment where (c) shows the area ratio (*A_ratio_* = *A_mt_/A_inter_*) and (d) shows the normalized roundness (*R_norm_*). (e-f) Contour plots for CCD experiment where (e) shows the area ratio (*A_ratio_*) and (f) shows the normalized roundness (*R_norm_*).

A region of parameter space was selected where the error in mitotic rounding measurements (*A_ratio_* and *R_norm_*) was minimized as shown in the Pareto front (Fig. 7b, Fig. 7e-f, SI-S9). The region of parameter space closest to experimental values of cell area and roundness was explored with a central composite design (CCD) as a second iteration to more precisely determine the relative contribution of each physical parameter on MR within experimentally observed ranges (Fig. 7a) [75]. This result quantitatively defines the predicted variation in mitotic cell-cell adhesion, stiffness and pressure that explains the variation in mitotic area ratio and rounding observed in mitotic epithelial cells. To keep mitotic rounding within the range of variation observed, AP must be tightly regulated (2D units of range are 19-23 nN/micron, ~19% variation in range about the calibrated point), whereas the requirements for 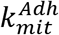 and 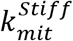 are less stringent ~120% and ~67% respectively.

Model reduction (SI-S10) revealed that regulation of mitotic rounding is approximated well by linear regression models for parameter evaluation resulting in physiological values of *A_ratio_* and *R_norm_* for *Drosophila* wing disc cells (Fig. 5). This suggests that regulation is in the linear regime, which is a good attribute for tightly controlled processes. Since interaction terms are not significant, cell mechanical properties effectively independently contribute to cell shape changes. Mitotic pressure was found to be the primary regulator of mitotic cell area (Fig. 7e), while both cell-cell adhesion and cortical stiffness reduced area expansion slightly. An increase in cell-cell adhesion was shown to reduce roundness whereas increased cortical stiffness promoted roundness for small perturbations (Fig 7f).

To define the relative impacts of mechanical properties on *A_ratio_* and *R_norm_* under physiological or “wild-type” conditions, local sensitivity analysis (Fig. 8a, b, SI-S11) was performed after application of the stepwise model reduction (SI-S10). Within the physiologically relevant domain of the parameter space, pressure strongly regulates mitotic area expansion but does not have a strong impact on the shape roundness (Fig. 7e, 8a). Stiffness and adhesion are important in tuning the degree of mitotic roundness (Fig. 7f, 8b).

**Fig. 8.**
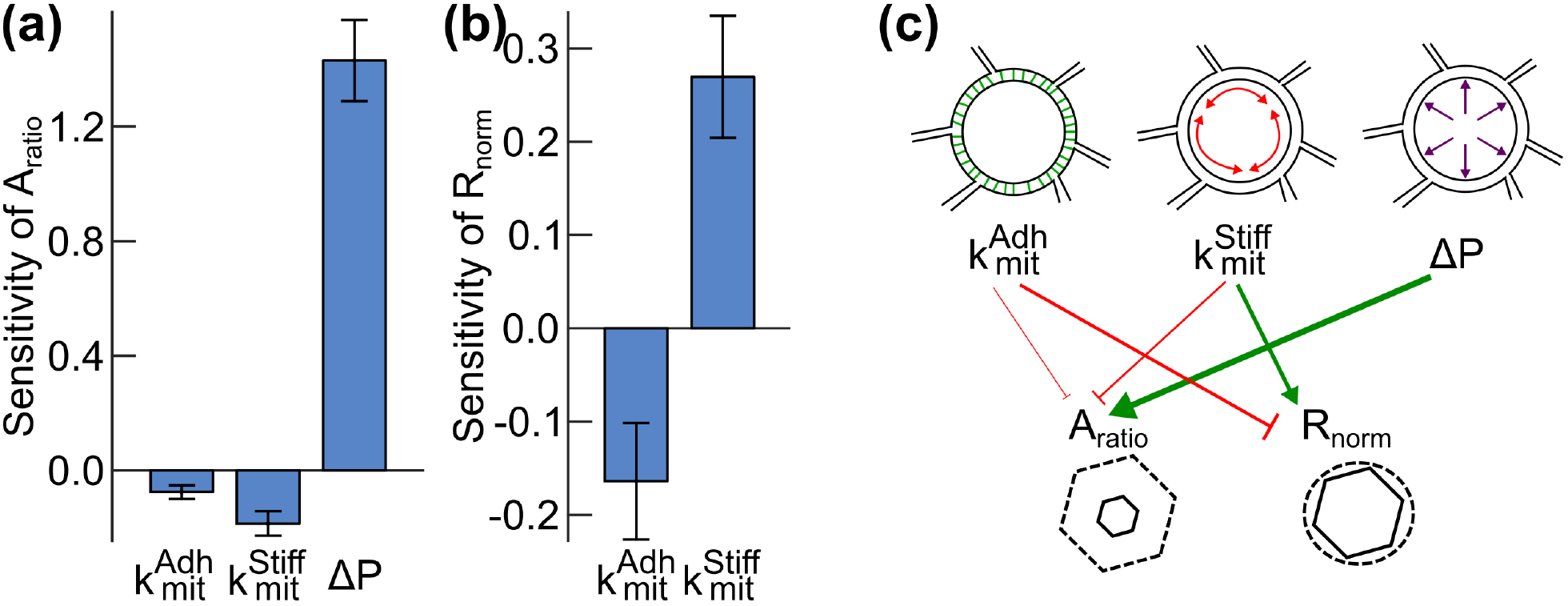
Quantitation of relative sensitivity of mitotic area expansion and roundness to adhesion, stiffness and pressure changes within the physiological property space. Sensitivity estimation of (*A_mit_/Ainter*) (a) and *R_norm_* (b) to small perturbation in the three mitotic parameter set points, 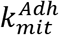, 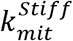 and Δ*P*. Sensitivity was estimated from the reduced RSM model described in Fig 7c-f after stepwise model regression (p-value cutoff of 0.01). (c) Proposed mechanical regulatory network defined for “physiological ranges” within the parameter ranges defined by the CCD (Run 2, Fig. 7a) that summarizes the local sensitivity analysis. Cell adhesivity, an increase in 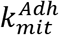 slightly inhibits area expansion and strongly inhibits roundness. Membrane stiffness, 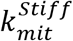 inhibits area expansion and promotes roundness. Mitotic area expansion is most sensitive to variation in the mitotic pressure change (ΔP), but pressure has little effect on roundness over the calibrated physiological ranges.

These results are summarized in the form of the mechanical sensitivity model in Fig. 8c, analogous to protein interaction networks. This model describes how small variations in each cellular mechanical property impact relative mitotic area expansion and roundness.

## Discussion

The roles of pressure, stiffness and adhesion in mitotic cells even in a single cell culture, in suspension or attached to substrates, are still not resolved in the experimental literature and largely unexplored in the tissue context [11,14]. We described in this paper a novel multi-scale sub-cellular model, called Epi-Scale, for simulating mechanical and adhesive properties of cells in the developing columnar epithelium of the wing disc, which consists of a single layer of cells. The model approximates the tissue as a 2D surface since the majority of the contractile and adhesive forces are localized at the apical surface of the epithelium (Fig. 1 b).

Parameter ranges for the computational model were obtained by calibrating the model using single cell stretching experiments, experiments on stretching a pair of cells adhered to each other, dynamic experimental measurements of the area and roundness of mitotic cells, and tissue growth rate of *the Drosophila* wing disc. Cell-cell adhesion and cell elasticity were calibrated using data from experiments with single cells. The calibrated model was verified by successfully reproducing emergent properties of developing tissue such as the polygon class distributions for both interphase and mitotic cells without additional calibration or parameter tuning.

Epi-Scale enables the systematic generation and testing of new hypotheses about the underlying mechanisms governing mitotic rounding within the developing tissue microenvironment. Regression analysis of predictive simulations provided complete assessment of the quantitative contributions of cytoplasmic pressure, cell-cell adhesion and cortical stiffness to mitotic cell rounding and expansion (Fig. 7, 8). Mitotic cell area expansion was shown to be largely driven by regulation of cytoplasmic pressure. Surprisingly, the variability in mitotic roundness within physiological ranges was shown to be primarily driven by varying cell-cell adhesivity and cortical stiffness, rather than pressure.

It is currently challenging to target only dividing cells in a tissue. One experimental approach that might be used in the future for testing the model predictions would be to regulate the expression of E-Cadherin, Myosin-II, and osmotic channel antagonists under a Cyclin B promotor, active during mitosis, resulting in modulation only in dividing cells [76,77]. Alternatively, opto-genetic methods could be employed to selectively regulate individual cell properties [78].

The simulation results have also shown that increases of the mitotic rounding under super-physiological pressure (greater than calibrated values) could result in cell-cell rearrangements (T1 transitions) of the neighboring cells, due to rapid increase of the apical surface of the mitotic cell (Fig. S9). This indicates that Epi-Scale platform could be used for future detailed studies of epithelial morphogenesis.

We have shown that our model simulations provide new insights into the individual contributions of cell properties to MR. Determining which aspects of mitotic rounding are most sensitive to perturbed cell properties in dense tissues, including solid tumors, can help direct future efforts to identify cellular processes that specifically block mitosis in highly proliferative tumors, but that are not damaging to non proliferative cells [14]. As a flexible computational modeling platform, Epi-Scale can be extended to simulate a wide range of multi-cellular processes, including epithelial morphogenesis, wounding healing and blood clot formation.

## Acknowledgements

MA, JZ, PB, CN, AN, AA were supported by the NSF Award CBET - 1403887. MA and ZX were also supported by the NIH grant 1U01HL116330. JZ was supported in part by CBET-1553826, PB was supported in part by Walther Cancer Foundation Interdisciplinary Interface Training Project, and CN was supported in part by the Notre Dame Advanced Diagnostics & Therapeutics Berry Fellowship (CN). The authors also acknowledge support from the Notre Dame Integrated Imaging Facility. They thank members of the Zartman’s lab for critical feedback on manuscript drafts as well as Luis Lazalde for technical analysis of segmented cells.

